# Static and Dynamic Functional Connectivity Analysis of Cerebrovascular Reactivity: An fMRI Study

**DOI:** 10.1101/580621

**Authors:** N. Lewis, H. Lu, P. Liu, X. Hou, E. Damaraju, Armin Iraji, V. Calhoun

## Abstract

The human brain, as a finely-tuned system, needs a constant flow of oxygen to function properly. To accomplish this, the cerebrovascular system ensures a steady stream of oxygenation to brain cells. One tool that the cerebrovascular system uses is cerebrovascular reactivity (CVR), which is the system’s ability to react to vasoactive stimuli. Understanding CVR can provide unique information about cerebrovascular diseases and general brain function. CVR can be evaluated by scanning subjects with blood-oxygenation-level-dependent (BOLD) functional magnetic resonance imaging (fMRI) while they periodically inhale room air and CO_2_-enriched gas, a powerful and widely-used vasodilator. Our goal is to understand the effect of vasodilation on individual intrinsic connectivity networks (ICNs), as well as how functional network connectivity (FNC) adapts to the same vasodilation. To achieve this goal, we first developed an innovative metric to measure the effect of CVR on ICNs, which contrasts to the commonly used voxel-wise CVR. Furthermore, for the first time, we studied static (sFNC) and dynamic (dFNC) FNC in the context of CVR. Our results show that network connectivity is generally weaker during vascular dilation, and these results are more pronounced in dFNC analysis. dFNC analysis reveals that participants did not return to the pre-CO_2_ inhalation state, suggesting that the one-minute period of room-air inhalation is not enough for the CO_2_ effect to fully dissipate in humans. Overall, we see new relationships between CVR and ICNs, as well as how FNC adapts to vascular system changes.

## INTRODUCTION

CVR reflects dilation and constriction capacity of blood vessels in response to different stimuli, particularly stress. Studying CVR provides an important perspective on how the brain functions in conjunction with the vascular system, which could lead to a greater understanding of cerebrovascular disease (Fierstra, Sobczyk et al. 2013). CVR can be measured by inducing vasodilation, such as inhalation of a CO_2_ gas mixture, while monitoring perfusion-sensitive MRI signals such as blood oxygenation level dependent (BOLD) MRI (Lu, Xu et al. 2011). These methods have been used in previous research to study cerebrovascular disease (Yezhuvath, Uh et al. 2012, Marshall, Lu et al. 2014) and to study brain networks related to CVR (Liu, Welch et al. 2016).

Currently, CVR is typically measured at a per-voxel level (Lu, Liu et al. 2014) by conducting a linear regression between voxel-wise time course of the BOLD signal and end-tidal (Et) CO_2_, which is the CO_2_ content in the exhaled air and an estimate of arterial CO_2_ level in an individual’s central nervous system. In this research, we expanded the concept of measuring CVR from the per-voxel level to a per-network level. The per-network CVR provides an understanding of pathophysiology as it relates to functional networks. The relationship between vasodilation and network level analysis could provide deeper insight into how between-network connectivity is altered, moving beyond spatial patterns to provide information about the ongoing dynamics. We accomplished this by calculating the correlation between the EtCO_2_ time courses and each network time course and then averaging this correlation across all subjects.

Our research further sought to develop a better understanding of how CVR and vasodilation relate to ICNs as well as the virtually unstudied area of how vasodilation due to CO_2_ inhalation impacts the connectivity relationships between ICNs. We accomplished this by observing and analyzing the influence of the CO_2_ inhalation effect on functional network connectivity (FNC), estimated as the cross-correlation between ICN time-courses via independent component analysis (ICA). We were also interested in evaluating how functional domains within the brain work together when reacting to cerebrovascular stress.

In order to study how the brain’s neural function changes due to vasodilation, 54 subjects were measured with BOLD scans during a CO_2_ task study. The subjects were asked to inhale a CO_2_ gas-mixture for approximately 60 seconds, and were then asked to breathe room-air for approximately 60 seconds. These intervals occurred three times over the course of the experiment. Once this data was collected, it was processed with group independent component analysis (gICA) to obtain a total of 100 brain ICNs (Calhoun, Adali et al. 2001, Himberg, Hyvarinen et al. 2004, Erhardt, Rachakonda et al. 2011), of which, 42 components were selected to best represent the brain networks. ICA is an effective tool as it is data-driven, and as such, preserves the vascular relationship within and between networks (Anderson, Druzgal et al. 2011). From this data, we segmented the data into CO_2_ and room-air groups based on the EtCO_2_ time-courses. To begin, we shifted the EtCO_2_ timecourses to align with the BOLD timecourses. The per-subject average of the EtCO_2_ timecourses was calculated and used as a threshold, such that all time points with a value below the average were considered to be periods of room-air inhalation, and all time points above were considered CO_2_ inhalation periods. Static FNC was applied to the CO_2_ and room-air segmented ICN timecourses in order to capture the overall changes of the FNC time course, while dynamic FNC was used to capture more nuanced changes that may have been lost while using the full time course to measure FNC. In order to include the time-varying information during the subjects’ transitions between the different air-composition intervals, we segmented the dFNC timecourses, as opposed to the ICN timecourses prior to dFNC. The results of both methods were captured and compared between the room-air and CO_2_ conditions.

## METHODS

### Data collection and Acquisition

The study was approved by the Institutional Review Board (IRB) of the University of Texas Southwestern Medical Center at Dallas. Each participant gave written informed consent. The dataset included 54 healthy participants that were part of the Dallas Lifespan Brain Study (DLBS) (Rodrigue, Rieck et al. 2013) and consisted of 22 males and 32 female young adults with ages ranging from 20 to 39. During the scans, each subject inhaled a CO_2_ gas mixture for approximately 60 seconds, and then spent another 60 seconds breathing room-air. This cycle recurred for a total of three times (Figure 1).

**Figure 1:**
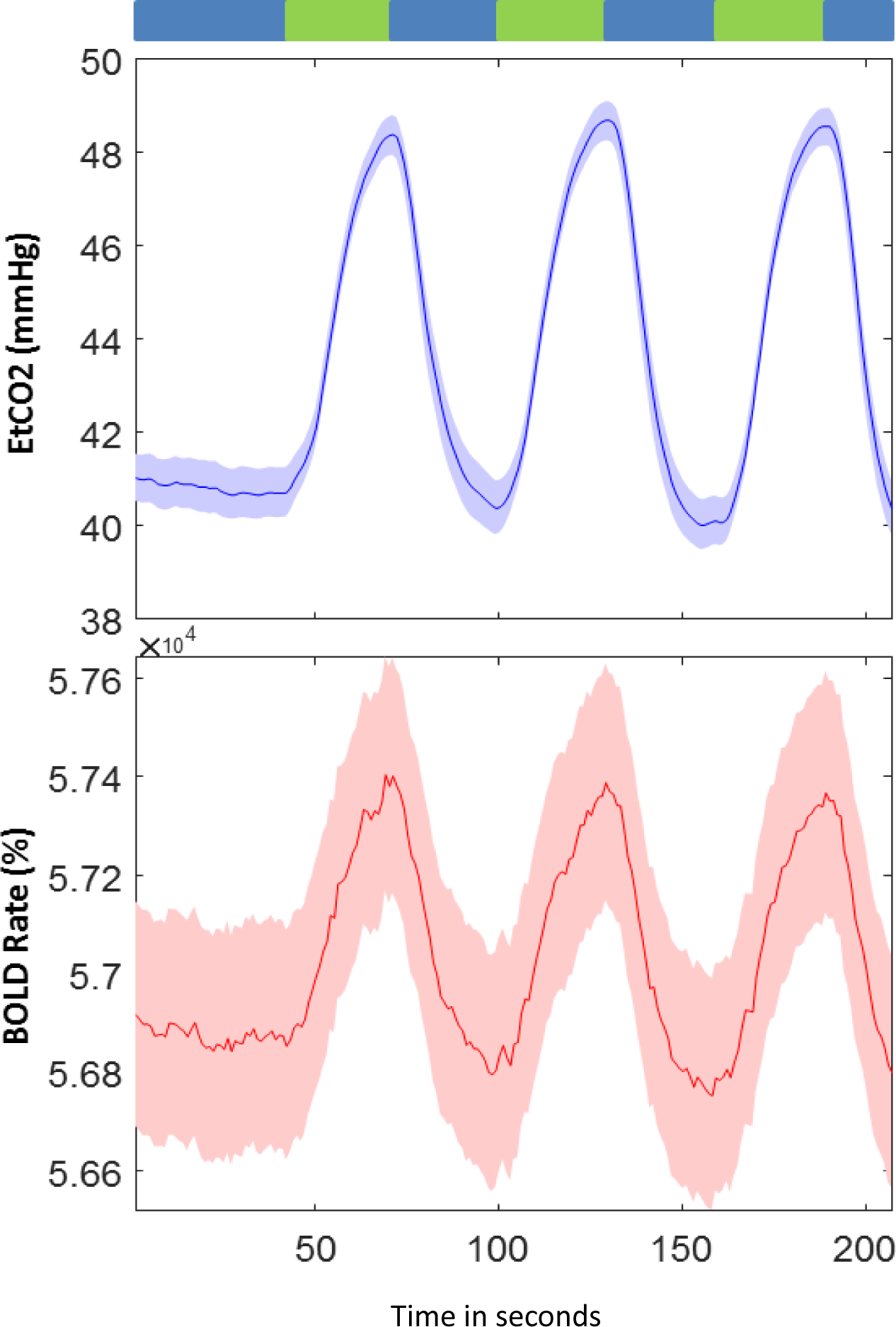
Mean and standard error across individuals of the paradigm of gas inhalation (top), and the concomitant CO_2_ (middle) modulation. As well as the mean and standard error across individuals of the BOLD signal time courses (bottom). The x-axis represents time in seconds.

### CO_2_ information

During the CVR scan, the subjects began by breathing normal room-air for 60 seconds (30 time points), and then breathed a gas-mixture with an elevated level of CO_2_ for another minute. From this point forward, the subjects cycled inhalation methods every 60 seconds (30 time points). The scan paradigm is shown in Figure 1, which shows the breathing task in detail. CO_2_ gas-mixture consisted of 21% O_2_, 74% N, and 5% CO_2_. This mixture has an elevated level of CO_2_, compared to the normal room-air, which is generally negligible, but with similar levels of O_2_ and N_2_ (Liu, Welch et al. 2016).

### Imaging parameters

The data were collected on a 3-Tesla Philips MR system with a 32 channel head coil. The BOLD fMRI images were acquired using an echo planar imaging (EPI) sequence with a repetition time (TR) of 2s, a field of view (FOV) of 220 × 220 × 150 mm^3^, voxel size of (3.4 mm, 3.4 mm, 3.5 mm), 43 total slices, a 64 × 64 matrix, and a total of 207 volumes. The echo time (TE) was 25 ms, and the flip angle was 80°.

### Data preprocessing

Preprocessing was performed primarily within the SPM software (http://www.fil.ion.ucl.ac.uk/spm/) and custom Matlab (https://www.mathworks.com) scripts. We used the INRIAlign toolbox in SPM to correct for subject head motion. Next, we performed a slice-timing correction using SPM. The data were then warped to the Montreal Neurological Institute (MNI) template and resampled to 3 mm^3^ isotropic voxels. Next, spatial smoothing with a 6 mm full width at half-maximum (FWHM = 6 × 6 × 6 mm^3^). Finally, the time course of each voxel was z-scored (normalized using the time course variance).

### Preprocessing and ICA

Spatial-group ICA was used to estimate a total of 100 components using the group ICA of fMRI (GIFT) Toolbox (http://mialab.mrn.org/software/gift/) (Calhoun, Adali et al. 2001, Calhoun and Adali 2012). Subject-specific principle component analysis (PCA) was first used to reduce the subject level time course to 200 principal components. Next, the principal components of individual subjects were temporally concatenated, and the group-level PCA was used to reduce the aggregated components from 200 to 100 along the direction of maximal variance across subjects (Erhardt, Rachakonda et al. 2011). The infomax algorithm was applied to maximize spatial independence of the group PCA reduced data resulting in a total of 100 components. The ICA algorithm was repeated 20 times and the most central run was selected to ensure stability (Ma, Correa et al. 2011). A group information guided ICA (GIG-ICA) approach from GIFT was used to back-reconstruct the subject specific spatial maps and time courses (TCs) from group-level independent components (Du and Fan 2013). The GIG-ICA approach has been shown to be a more effective artifact removal approach than using single subject ICA prior to the group ICA analysis (Du, Allen et al. 2016) and more sensitive to group differences than a spatiotemporal regression approach (Salman, Du et al. 2017).

### Post ICA processing

ICNs were identified by assessing the spatial maps and time courses of the independent components. One sample t-tests for each spatial map were calculated and then those maps were thresholded to obtain the peak activations for each component. ICNs were selected if the peak activations covered gray matter and showed minimal overlap with vascular, ventricular, or edge regions (Allen, Erhardt et al. 2011). The mean spectral power (Allen, Erhardt et al. 2011) was calculated for the TC corresponding to a given component. This information, along with a priori knowledge of ICNs was used to select ICNs. These methods resulted in a total of 42 ICNs. We then associated each ICN with a specific functional domain, based on our prior work (Allen E.A. 2012). The seven domains consisted of: subcortical (SC), auditory (AUD), sensory-motor (SM), default mode (DM), Attention (ATN), Visual (VIS), and Cerebellar (CB). Once the ICNs were selected, the corresponding TCs were detrended and then despiked for FNC analysis. As it has been shown to be effective in reducing noise, a low-pass band filter (0.15 Hz) was used to preprocess the ICN time courses prior to computing the FNC (Allen, Erhardt et al. 2011).

### Data Partitioning

The EtCO_2_ time courses were used to determine in which time points the subjects were exposed to either the CO_2_ gas-mixture or room-air. The subject-wise time course was thresholded as either below the average or above the average, defining the room-air and CO_2_ conditions used in our experiments. To mitigate noise associated with ambiguous time points, or those steps in which the subjects were transitioning between intervals, we also experimented with groups where scans at the beginning and end of each CO_2_ and room-air interval were eliminated. The results from these comparisons showed no statistically significant differences from one another and thus we report only on the results using the above/below mean approach. However, the time points at the beginning and end of each interval might be ambiguous due to the subjects’ transition to or from CO_2_ inhalation. In order to evaluate the impact of this, we performed the same experiments while removing the first and last time point of each interval. Our results and conclusions were effectively the same as with the case in which all time points were used. The portioning was done after ICA, but the order with respect to the FNC calculations differed based on whether the FNC was static or dynamic. In the case of sFNC, the subject-wise BOLD time courses were separated before the FNC matrices for both groups were individually calculated. In the case of dFNC, the FNC timecourses were portioned into the two groups. The act of portioning the timecourses after the dFNC calculations was done so as to not bias the FNC timecourses by group.

### Network-Wise CVR Calculation

As this work focuses primarily on functional network analysis, it was pertinent to quantify the effect of CO_2_ at the network level. This informs us as to which networks are most impacted by vascular reactivity. In order to approximate the network-wise CVR, we calculated the regression coefficients between each network time course and the EtCO_2_ time course for every subject. These coefficients were then averaged across all subjects per network and weighted with the network spatial maps to better visualize network-wise CVR. Because the ICN timcourses were z-scored, the standard deviation of the timecourses is 1. As such, the correlation values are the same as CVR calculations except for a global scaling value, the standard deviation of the EtCO_2_ timecourses. Due to this similarity, we can rationally use correlation to represent the effect of CVR on the ICNs. We do this as correlation demonstrates the strength of the similarity between the EtCO_2_ and ICN timecourses.

### Static Functional Network Connectivity (sFNC)

To compute sFNC, the timecourses were segmented into either room-air or CO_2_ intervals based on the EtCO_2_ average for every subject. Then, the pairwise correlations between ICN time courses were calculated for each subject, which results in a 42 by 42 symmetric FNC matrix. The columns and rows of the correlation matrix were ordered by the aforementioned domains.

### Dynamic Functional Network Connectivity (dFNC)

In addition, a dynamic FNC analysis was performed on the entire component time courses, including both room-air and CO_2_ time points, which includes sliding-window correlation followed by clustering (Allen E.A. 2012). The chosen window size was 30 TR (60 seconds) in steps of 1 TR, consistent with previous work suggesting this is a good trade-off between over smoothing and sensitivity to noise (Vergara and Calhoun 2018). To allow for tapering along the edges, each window was defined as a rectangular window of 30 time points, convolved with Gaussian with a 3 TR full width at half max. We estimated covariance from the regularized inverse covariance matrix (ICOV) using the graphical LASSO framework to reduce noise associated with short time series (Allen, Erhardt et al. 2011). In order to impose sparsity, we imposed an L1 norm constraint on the inverse covariance matrix. The log-likelihood of unseen data was evaluated to optimize the regularization parameter for each subject in a cross-validation framework. The dFNC timecourses or the correlation matrices per time point for every subject were then segmented based on the EtCO_2_ thresholding method.

### Clustering

As has been observed in the past, patterns of network connectivity can reoccur within subjects across time and across subjects. Because of this, we used k-means to cluster the FNC windows in order to minimize the distance between members of a cluster and its cluster centroid (Allen E.A. 2012). We used the city-block distance as our measure, due to previous research that suggested city-block was more effective than Euclidean (Aggarwal C, Hinneburg A et al. 2001). The elbow criterion was used to estimate the model order of five clusters. Initially, we clustered a subset of windows (known as subject exemplars) from every subject corresponding to the windows with maximal variance in correlations between component pairs. The exemplars were obtained by calculating the variance in connectivity across all ICN pairs at each window and selecting windows corresponding to local maxima among this variance time course. From this, we clustered the exemplars and calculated the 5 centroids. These centroids were then used to initialize a clustering of the entire dataset.

### Statistics

From the sFNC results, we computed average matrices across all subjects for both the CO_2_ and room-air results. We also computed a paired t-test per ICN pair, for CO_2_ versus room-air. The same method of comparison was used for each of the five dFNC states between the CO_2_ and room-air matrices. From the dFNC matrices, we also calculated several additional analyses. We computed the transition matrices, or the probability of a subject changing from one state to another, between the five states for both the CO_2_ and the room-air results, and then compared the two transition matrices with a paired t-test. The mean dwell time (MDT), or how long a subject stayed in a single state without changing states, and the fraction rate (FR), or how often a given state occurred, were also calculated for both CO_2_ and room-air results and then compared via a paired t-test.

## RESULTS

Building upon previous whole-brain functional connectivity work, we estimated and evaluated ICNs and their corresponding time courses using spatial group ICA. From the data we were able to categorize a total of 42 ICNs using our previous knowledge of brain anatomy. As mentioned, we categorized the ICNs into the seven aforementioned domains. Using these networks, we calculated the network-wise CVR. We then examined the static functional network connectivity (sFNC), or the temporal correlations between ICNs across the entire time course (Jafri, Pearlson et al. 2008). We also investigated the dynamic functional network connectivity (dFNC), or the functional network connectivity with a sliding-window approach (Allen E.A. 2012). Both sFNC and dFNC were performed separately for the CO_2_ and room-air time points, and then compared using a paired t-test.

### Network-wise CVR Calculations

We applied the network-wise CVR measurement technique on this data, the results of which can be seen in Fig. 2. From these results, we can see the difference between voxel-wise CVR and network-wise CVR. The voxel-wise CVR tends to be more prominent in the grey-matter regions of the brain, as that is where much of the brain’s vasculature resides. The network-wise CVR, although it does show similarities to the voxel-wise CVR, there were key differences within certain parts of the brain. There are clear areas of low correlation which can be observed in the network-wise CVR maps. This would appear to be caused by a lack of individual networks in those areas. However, there were also regions that were lower in the voxel-wise maps, possibly due to increased noise in the voxel-wise measurements and the multivariate nature of the network-wise CVR maps. Notably, there were prominent networks with low correlation to the CO_2_ effect, including components 79 and 55 in the DMN domain, as well as network 9 in the SC domain. Aside from the differences, the highest and most consistent correlations occurred in networks 18, 19, and 23 in the VIS domain, and network 54 in the DMN domains respectively. There were also several SM networks with relatively high correlation to the CO_2_ effect. These network names can be identified in figure 2.

**Figure 2:**
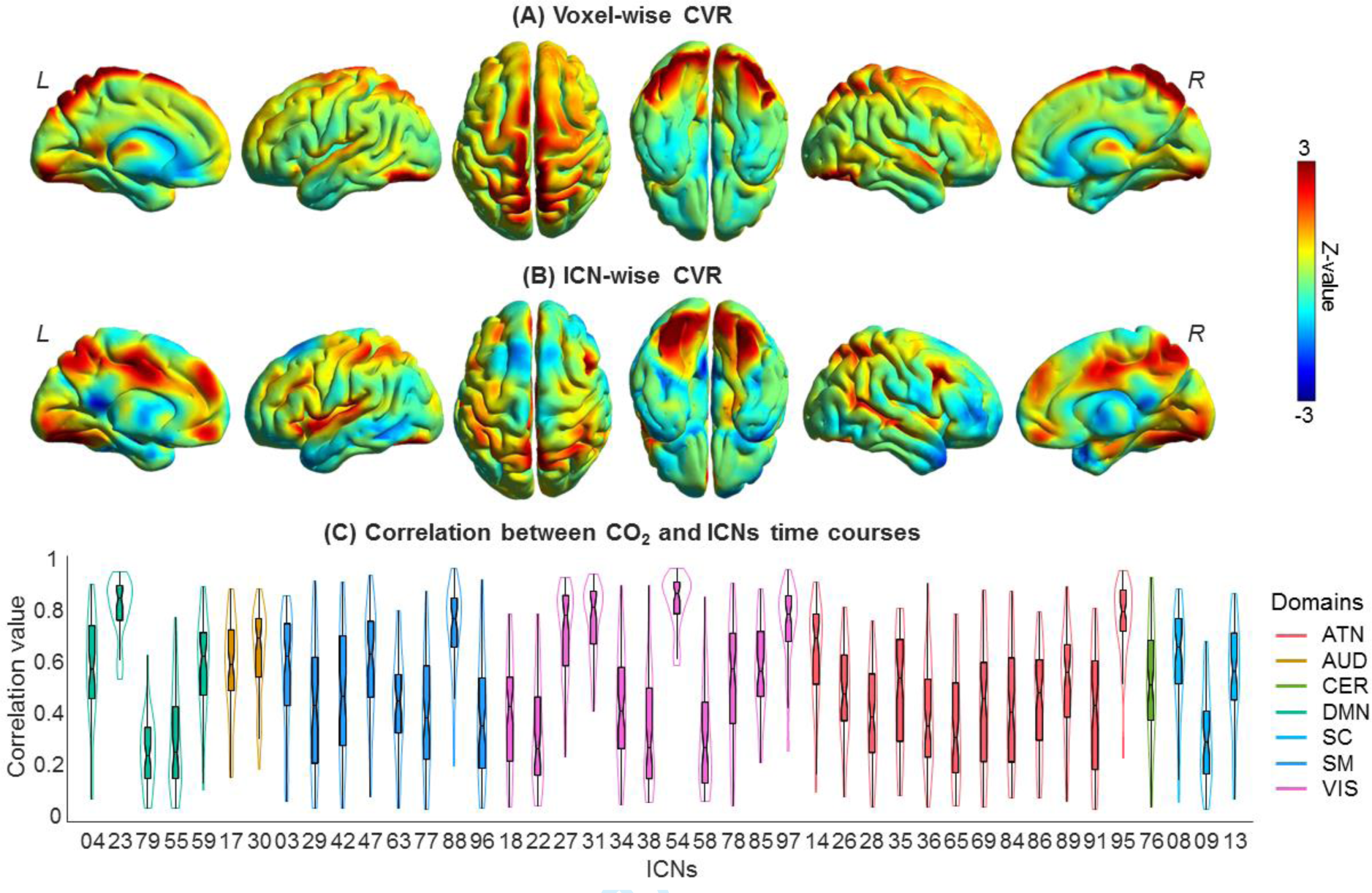
A) The voxel-wise CVR map compared to B) the network-wise CVR map, showing (from left to right), interior left hemisphere, exterior left hemisphere, top, bottom, exterior right hemisphere, and interior right hemisphere. C) A violin plot showing the median value, the interquartile range, the probability density, as well as the confidence (95%) interval of the network-wise CC values.

### sFNC results

The sFNC correlation matrices were calculated separately for CO_2_ and room-air time points. The paired t-test results provided a comparison between the CO_2_ and room-air matrices across all subjects. The resulting matrix (figure 4) was corrected for multiple comparisons using the false discovery rate (FDR), thresholded at 0.05.

**Figure 3:**
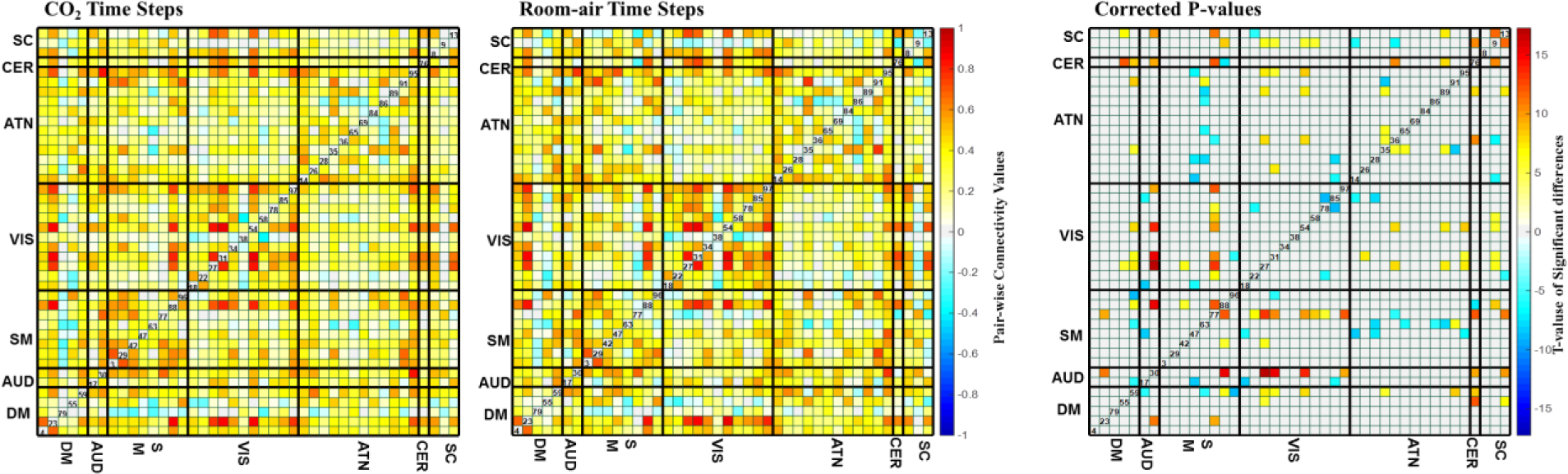
The mean sFNC maps for room-air (left) and CO_2_ (right) time points. The black lines separate the FNC maps into the seven domains, labeled as: subcortical (SC), auditory (AUD), sensory-motor (SM), default-mode network (DMN), Attention (ATN), Visual (VIS), and Cerebellar (CB). The group differences (using paired t-tests) between room-air and CO_2_ time points. These values are the FDR-corrected negative log of the p-values multiplied by the sign of the t-statistic. These values have been corrected for multiple comparisons via a false discovery rate (FDR) threshold of 0.05.

**Figure 4:**
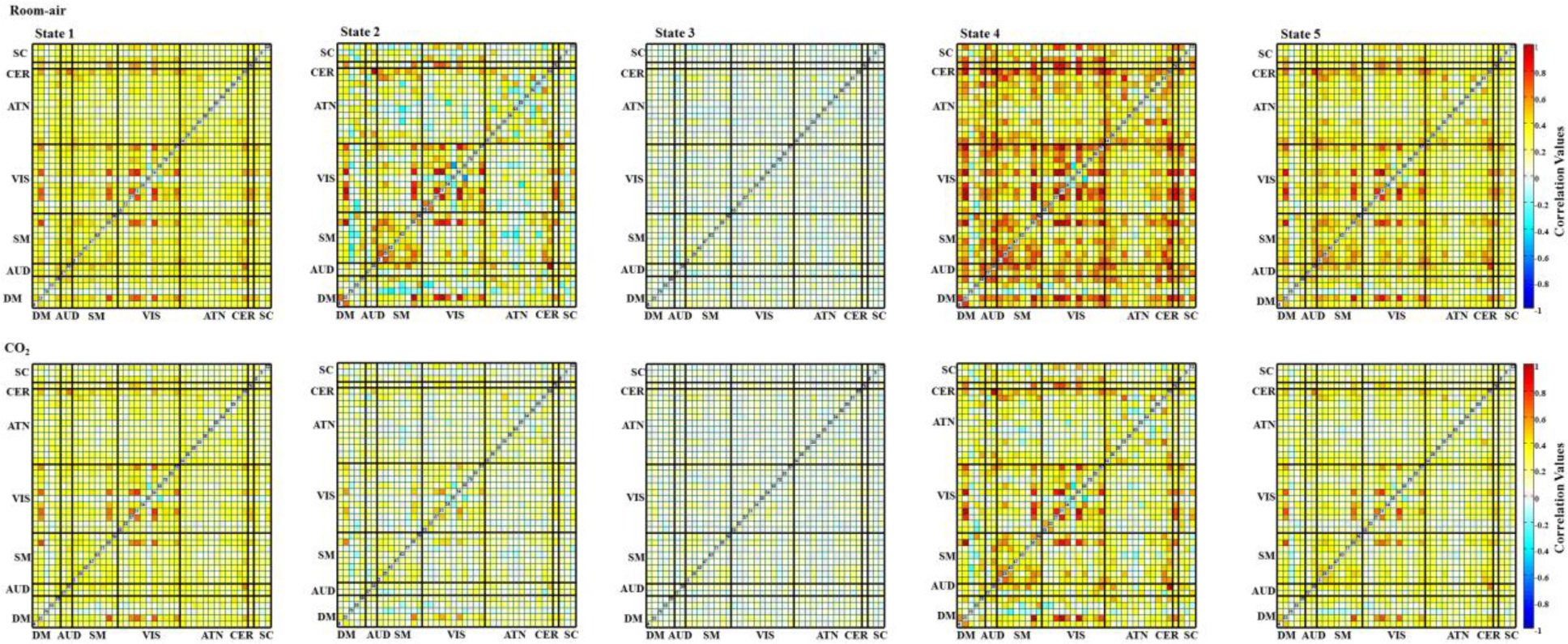
The mean dynamic FNC maps for both room-air (top row) and CO_2_ (bottom row) time points. Each column represents the FNC for each state, from state 1 to state 5. Significant cell-wide differences are visible in States 2, 4, and 5.

### dFNC results

States 2, 4, and 5 showed higher brain connectivity in the room-air time points, as opposed to the CO_2_ time points. This is expected since, as the oxygenation of the brain increases (due to venous oxygenation), the BOLD signal becomes less sensitive to oxygenation effects caused by neural activity (Boynton, Engel et al. 1996). Both room-air and CO_2_ time points showed higher connectivity (compared to other pair-wise correlations) within the VIS domain compared to other domain pairs (figure 5).

**Figure 5:**
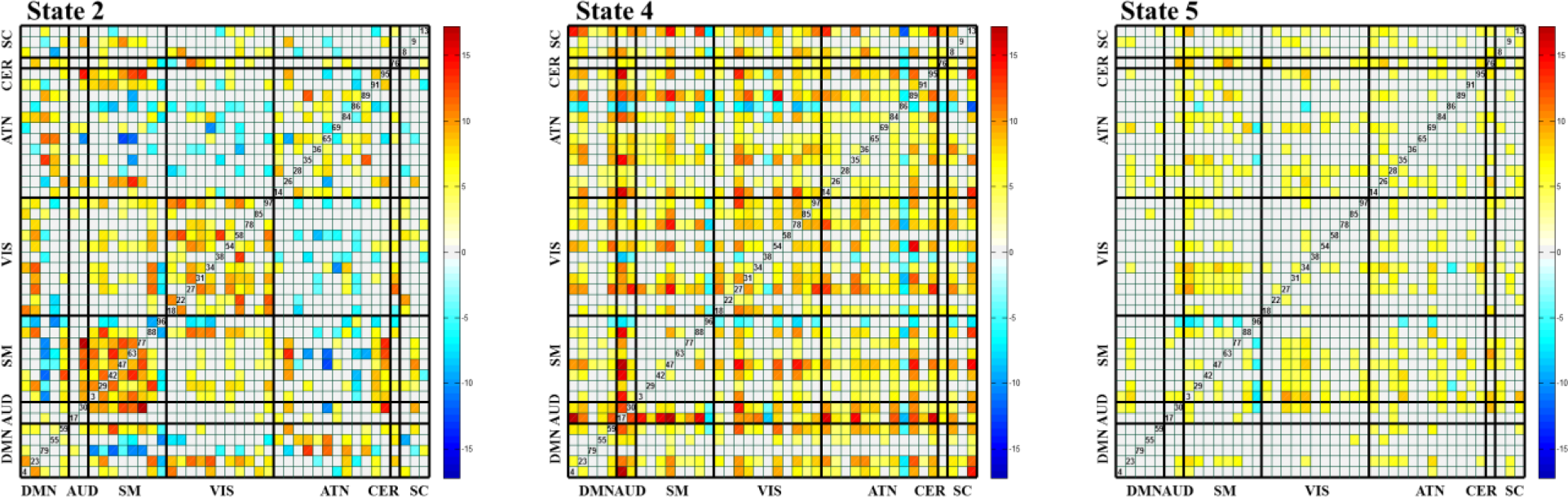
Paired t-tests for each pair-wise correlation of the dFNC maps for (left to right) states 2, 4, and 5 for room-air – CO_2_. The t-tests are the negative log of the p-values, corrected with a false discovery rate (FDR) threshold of 0.05, and multiplied by the sign of the t-statistic. States 1 and 3 were omitted because they had non-significant differences.

The five dFNC states, computed across all time steps regardless of CO_2_ content, were compared with paired t-tests across all subjects for each pair-wise correlation. From this, we saw the greatest differences in states 2, 4, and 5. The results can be seen in fig. 6. From this, we can see large differences between room-air and CO_2_ time points within the SM and VIS domains in state 2. The SM domain showed higher correlation overall. We also see smaller differences in the SM domain in state 4. Additionally, we see that state 2 most often occurred in the first portion of the experiment, with very little occurrence during the remainder of the experiment.

**Figure 6:**
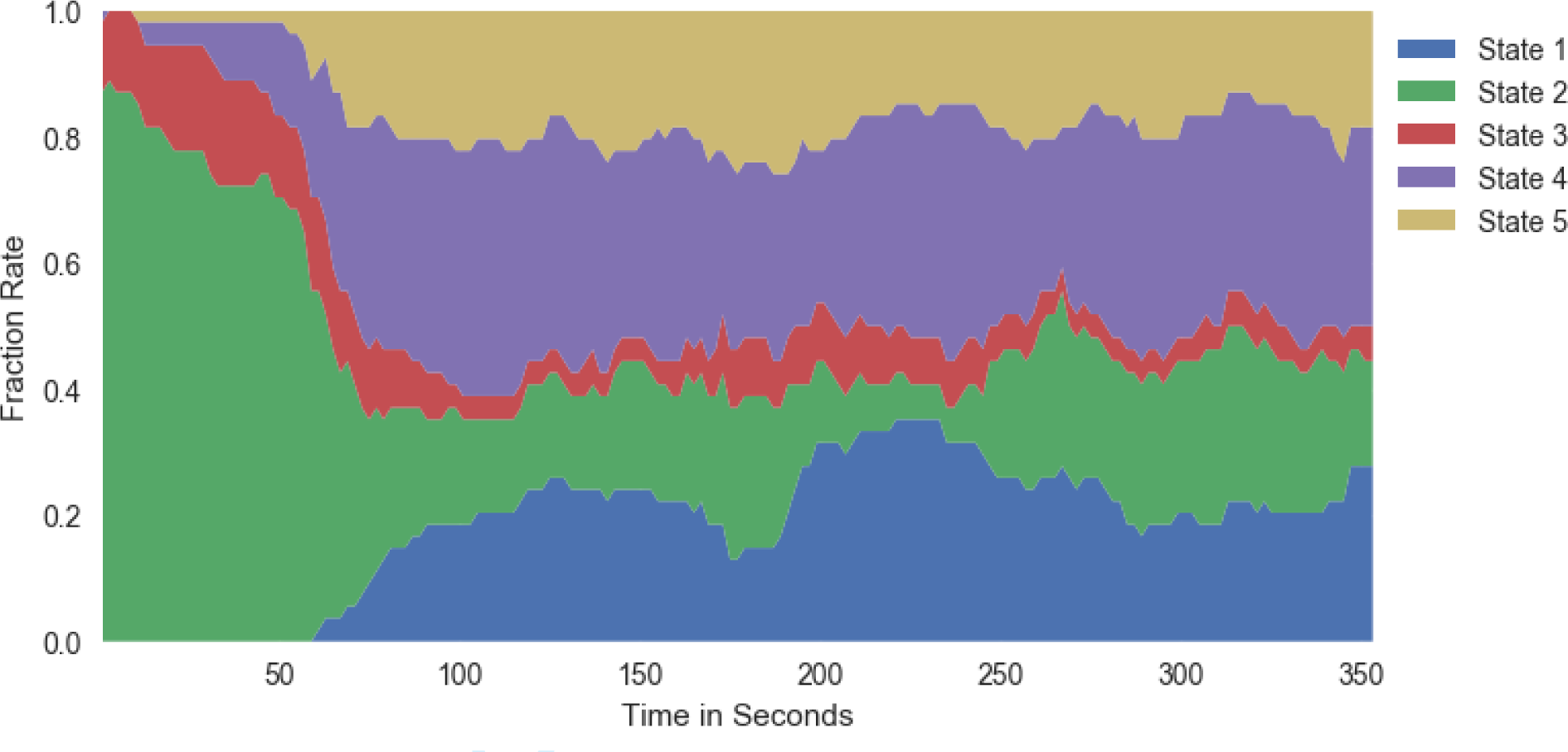
The percent occurrence of each state across the entire time series (averaged across all subjects). We see that state 2 occurs most often in the first portion of the experiment, before subjects began inhaling the CO_2_.

Figure 7 shows the per-time point occurrence of each state averaged across all subjects. This allows us to visualize changes in the dynamic connectivity which is consistent across individuals. We see that state 2 primarily occurs in the first segment of the experiment, before the subjects have inhaled any amount of the CO_2_ gas mixture. This aligns with our FNC results showing that state 2 had a significant difference between room-air and CO_2_ time point.

**Figure 7:**
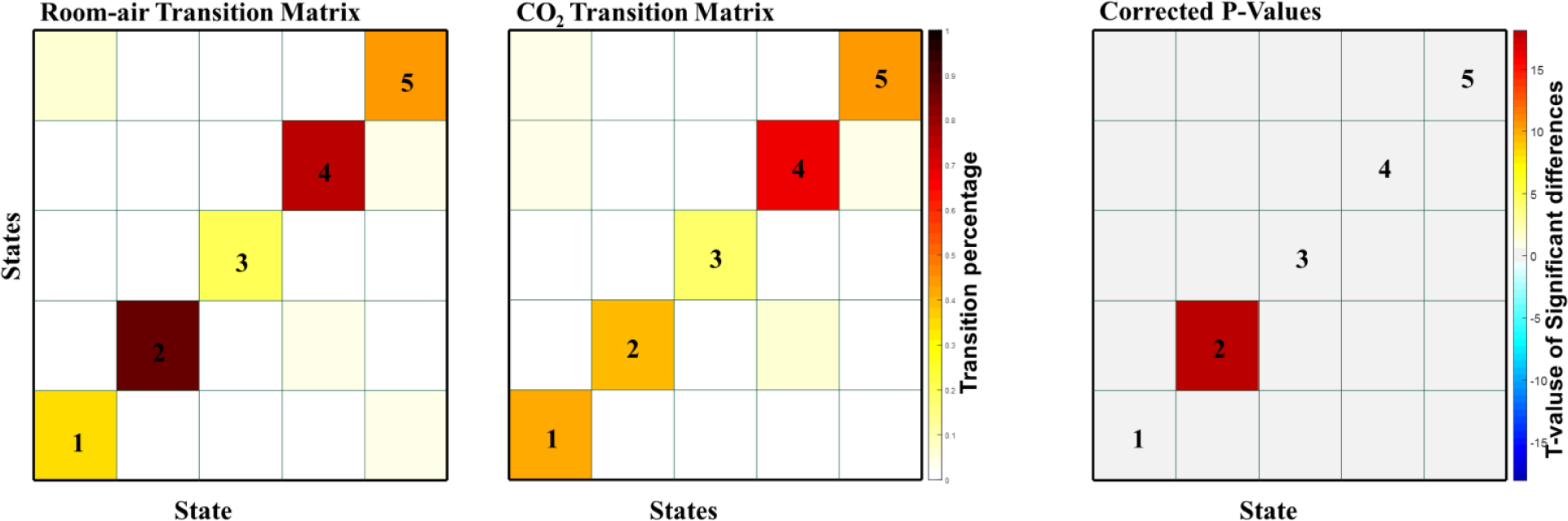
The transition matrices between all 5 states for room-air (left) and CO_2_ (middle) time points. The paired t-tests between CO_2_ and room-air time points (right) are the negative log of the p-values, corrected with an FDR correction with a threshold of 0.05. Only one transition cell, 2-2 passed the FDR threshold.

### Transition Matrices

After the dFNC states were calculated, we measured the transition probabilities between states. The transition matrices for both room-air and CO_2_ show high transition probability within states (fig. 8) and relatively low transition probabilities between states. FDR corrected paired t-tests show little difference between states room-air and CO_2_ transition probabilities, except for within state 2, which aligns with the other results related to state 2.

**Figure 8:**
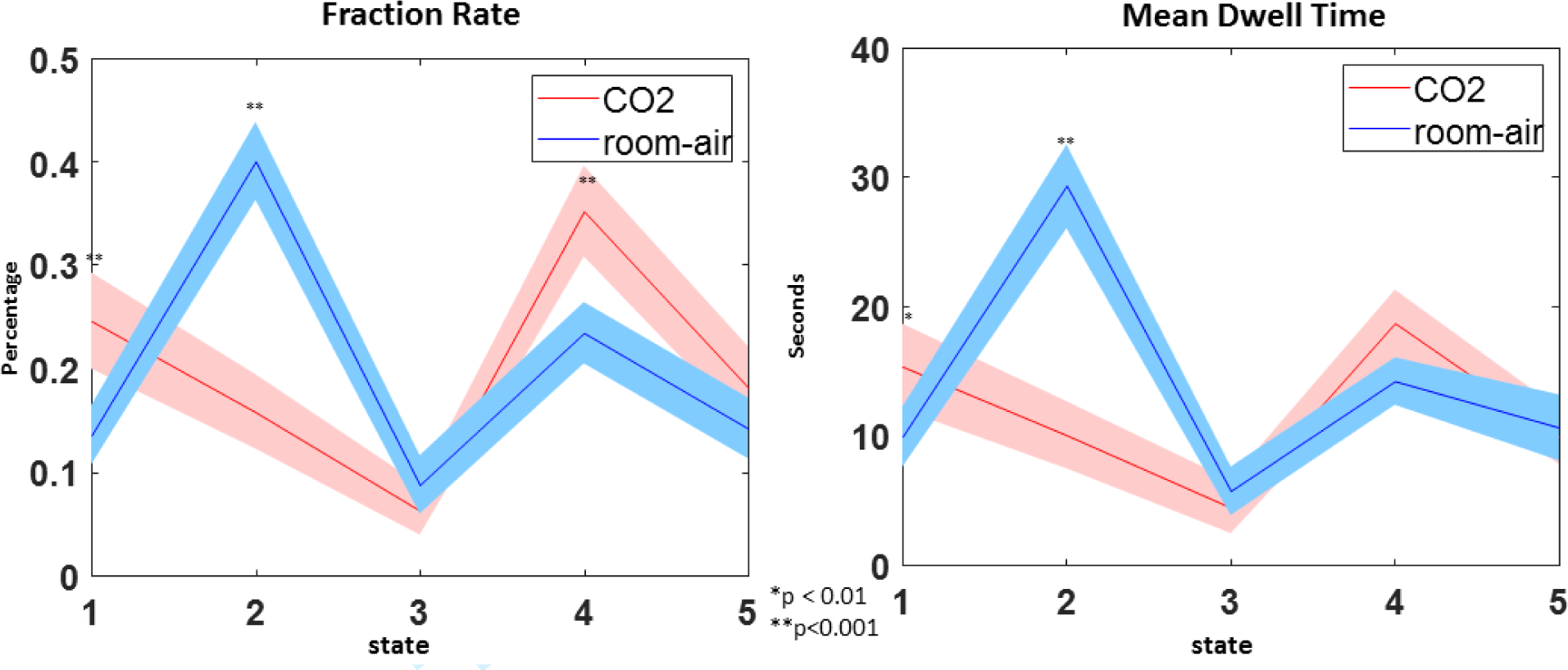
(left) The FR of all 5 states comparing CO_2_ and room-air time points. (right) Mean standard error of dwell times for all five states comparing CO_2_ and room-air time points.

### Mean Dwell Time Results

The mean dwell time (MDT), or the average number of consecutive time points a subject is classified as a given state, was also calculated for both room-air and CO_2_ time points. A paired t-test was used to compare the MDT per each state between the room-air and CO_2_ time points. MDT across all subjects showed significant differences between room-air and CO_2_ for states 1 and 2 (Fig. 7).

### Fraction Rate Results

The fraction rate (FR), or the total number of time points a subject is classified as a given state, were also calculated. These results can be seen in figure 9. As with the MDT results, paired t-tests were used to compare the FR of each state between room-air and CO_2_ time points. Similar to the MDT results, the FR of states 1, 2, and 4 were significantly different between CO_2_ and room-air time points.

## DISCUSSION

CVR is a powerful approach to study the human brain. In some recent studies it has been shown that large scale resting networks can be estimated from CVR data (Liu, Welch et al. 2016, Hou, Liu et al. 2019). However CVR data has not yet been studied in the context of high-order group ICA, nor have between network connectivity and associated dynamics been previously studied. Our analysis quantified the effect CO_2_ has on both sFNC and dFNC. This provides, for the first time, an evaluation of the connectivity across the entire brain during a CVR experiment. We showed, primarily from the dFNC results, that inhaling CO_2_ reduces the overall functional network connectivity. These results reinforce and extend previous research (Xu, Uh et al. 2011, Madjar, Gauthiera et al. 2012). Reduced functional connectivity can be seen both between domain and within domain.

We present novel results showing an estimation of CVR at the network level. This was accomplished by calculating the correlation between each network and EtCO_2_ timecourse, an estimation of CVR. The EtCO_2_ timecourse is used as an approximation of the CO_2_ content in arterial blood within the brain (Lu, Liu et al. 2014), which acts as the causal agent of vascular reactivity. A high correlation between the EtCO_2_ and network BOLD timecourses implies that vascular reserve in a given network is abundant. Results showed high correlation between the EtCO_2_ timecourse and BOLD signal in both the VIS and SM domains. A benefit of looking at the network-wise CVR relationship is that we can see the interplay between CVR and brain function. This interplay could be explored more in-depth in the future with a more nuanced analysis of the correlation between functional domains and CVR estimation. Future research could also focus on new techniques to evaluation the relationship between CVR and functional networks that are more nuanced than correlation alone.

Compared to the sFNC results, the dFNC results showed much larger differences between the room-air and CO_2_ results, both in local patterns and across the whole brain. This dynamic analysis provides nuanced information about how vasodilation impacts brain connectivity that is not detected in a static analysis. For example, the dFNC analysis captured information about how the brains changed from the beginning of the experiment (before CO_2_ inhalation) and the time points during the experiment. The dFNC analysis also appears to be more sensitive to the CO_2_-induced brain activity changes, capturing multiple significant changes in more than one state.

From the dFNC maps, as well as the state occupancy rates (Fig. 7), we consider state 2 to be a state in which the room-air time points are mostly free from the CO_2_ effect. We observed that state 2 has the most dominant anti-correlation patterns between functional domains, including patterns between the SM and DM domains, the SM and ATN domains, and between the VIS and ATN domains; patterns that have been seen in previous research (Fox, Snyder et al. 2005, Fox, Zhang et al. 2009, Uddin, Clare Kelly et al. 2009). These anti-correlation patterns are most dominant in state 2. Within this state, the CO_2_ effect had the highest impact on lowering the connectivity within the sensory-motor domain (seen in Fig. 6). This result is consistent with previous findings showing that CO_2_ impacts the SM domain (Liu, Hebrank et al. 2013, Mazerolle, Ma et al. 2016, Golestani, Kwinta et al. 2017), as well as the VIS domain, meaning that our findings that CO_2_ inhalation reduces connectivity within these two domains accompanies an overall reduction of activity within the two domains. These results are congruent with results from the network-wise CVR estimation as there was both a significant impact of CO_2_ on the FNC of SM and VIS domains, and both domains were highly correlated with the EtCO_2_ time courses. We suggest that this congruency adds robustness to our conclusions about both FNC differences and the effectiveness of our network-wise CVR estimation.

Based on our findings that there are marked differences between state 2 and the other states, we show that generally, the BOLD signal after the start of the CO_2_ inhalation was affected by the CO_2_ inhalation, even during the room-air intervals. This suggests to us that the minute-long periods of room-air inhalation may not be enough time for the average subject to recover from the neural modulation effects of CO_2_ inhalation.

State 4 also produced interesting results, in that it showed more global differences than the other states (fig. 6). This indicates that although CO_2_ has significant impact on specific regions, it also has a significant global effect as well. This result may indicate that the vasodilation caused by CO_2_ occurs indiscriminately across the entire brain. State 5 shows a similarly global difference, but with a smaller effect.

The MDT and FR are secondary metrics used to evaluate time-varying information of the FNC patterns, which gives us a broader perspective than what FNC maps alone provide. From our results, the FR and MDT show significant differences within state 2 (room-air > CO_2_), which is congruent with the FNC maps, but they also show significant differences within state 1 (CO_2_ < room-air). From Fig. 9, we speculate that this might be related to the fact that state 1 contains few time points from the initial portion of the experiment. The dFNC matrices show insignificant cell-wise differences within state 1, which implies that both room-air and CO_2_ intervals are impacted by the CO_2_ effect, as most of the time points occur after or during CO_2_ inhalation. However, the FR and MDT do show significant differences, which may be due to the lack of room-air time points clustered as state 1, meaning there are fewer room-air time points compared to CO_2_ time points. This could possibly increase the difference within the MDT between room-air and CO_2_ time points, and would definitely increase the FR differences. We also see that there are significant differences within the state 5 FNC maps between room-air and CO_2_, but no significant difference in the FR or MDT. It is also possible that this is, in part, due to the opposite effect found in state 1. Approximately 1% of all time points within the first interval are clustered as state 5. This may be the cause of some of the differences between the FNC maps, but may also contribute to the similarities found in the FR and MDT. The first interval is a slightly larger length of time than the other intervals, 80 seconds compare to 60 seconds. But, due to the lower number of state 5 time points within the first interval compared to the other intervals, there would be a more equal number of room-air time points and CO_2_ time points. This would show the opposite effect from state 1, as it may reduce the MDT differences, and would most likely reduce the FR differences.

## CONCLUSION

Our network-wise CVR calculation is a simple method to depict the relationship between CVR and individual ICNs. We suggest that this method could be used in future analyses of CVR to capture more of the spatial relationships using the multivariate connectivity networks. The network-wise CVR maps showed the relationship between CVR and networks, which, when compared to per-voxel CVR maps, show distinctive differences. We showed high network-wise CVR in both the SM and VIS domains.

The results from our experiments showed wide-spread differences between room-air and CO_2_ time points. We saw that across the whole brain, CO_2_ time points showed lower correlation values than the room-air time points. The dFNC analysis, likely a more natural way to analyze brain connectivity, shows more sensitivity to CO_2_ effects not detected by the sFNC. For instance, from the dFNC results, we concluded that state 2 was most prevalent prior to exposure to CO_2_. Given this, it may be useful to utilize this connectivity pattern to predict breathing normally or breathing a CO_2_ heavy air mixture. Due to the differences between the states, we also suggest that the minute-long period of time between CO_2_ inhalation intervals was not enough time for the subjects to fully recover from the CO_2_ gas-induced neural modulation. We also concluded that CO_2_ reduces correlation value across the whole brain for both global and local effects, with local effects primarily affecting the SM and VIS domains. The observed effects of CO_2_ on the SM and VIS domains are comparable with the network-wise CVR calculations which showed high correlation between BOLD signals in these domains and the EtCO_2_ time courses respectively.

## Acknowledgements

DLBS grants R37 AG006265 and R21 AG034318.

